# Clonal expansion of personalized anti-tumor T cells from circulation using tumor organoid-immune co-cultures

**DOI:** 10.1101/2020.11.06.371807

**Authors:** Qingda Meng, Shanshan Xie, G. Kenneth Gray, Weilin Li, Ling Huang, Dipikaa Akshinthala, Mohammad H. Dezfulian, Elizabeth Ferrer, Catherine Conahan, Sofia Perea Del Pino, Joseph Grossman, Stephen Elledge, Manuel Hidalgo, Senthil K Muthuswamy

## Abstract

Tumor-specific cytotoxic T cells are effective tools for cancer immunotherapy, but the ability to generate them continues to be a challenge. Furthermore, there are no compelling approaches to empirically identify tumor-targeting T cells and T cell receptors by exploiting the multitude of antigens on tumor cell surfaces. Here, we use patients’ peripheral blood and autologous tumor organoids to enrich tumor-specific cytotoxic T cells with patient-specific killing mechanisms and a tissue-resident memory phenotype. We further demonstrate that these organoid-primed T (opT) cells undergo several orders of magnitude of clonal expansion and express T cells receptors and check-point proteins unique to each patient. Importantly, transferring the TCRs to heterologous T cells was sufficient to confer tumor recognition in a patient-specific manner. Thus, we report a patient-specific and antigen-agnostic platform for expansion of tumor-targeting T cells and identification of cancer-targeting TCRs from the peripheral blood of pancreatic cancer patients that can be exploited for immunotherapy applications.

---

Identifying T cells with specific reactivity against human tumor cells would represent a significant advance for cancer immunotherapy. The tumor-selective T cells will also assist in empirically defining the antigenicity of cancer cells and in identifying T cell receptors (TCR) that can be utilized to engineer cell therapy products. Most of the efforts to identify tumor-targeting T cells are focused on the acquisition and expansion of tumor-infiltrating T lymphocytes (TILs). TILs, especially cytotoxic CD8+ T cells, are being used for adoptive immunotherapy with encouraging clinical results (*1–4*). Analysis of the TCRs in populations of TILs shows a broad range of clonal expansion with specific clones representing 0.1-50% (*5, 6*). However, the apparent lack of TIL populations in many carcinomas, including that of the prostate, ER+ breast, and pancreas and the intra-tumor heterogeneity for TIL populations (*7–9*), creates challenges for obtaining tumor-targeting T cell populations for broad cancer treatment. Furthermore, TILs are frequently associated with T cell exhaustion (*10–12*), which impairs their effectiveness when used for adoptive cell therapy (ACT).

In contrast to TILs, the TCR repertoire in the blood is more diverse with a wide range of epitope specificities. Identification of tumor-selective cytotoxic T cells from peripheral blood is typically performed by culturing T cells in the presence of tumor-associated antigens or antibodies targeting activated T cell markers such as 41BB and PD1 (*13–15*). Although effective, these approaches do not exploit the diversity of tumor cell surface because the expansion process is usually targeted towards known tumor-associated antigens. Recent studies suggest that peripheral blood can be combined with autologous tumor cells or 3D tumor organoids from lung cancer, melanoma, or pancreatic cancer (*16–18*) to generate populations of T cells with cytotoxic properties. However, whether tumor-targeting T cells obtained from peripheral blood undergo clonal expansion in response to exposure to tumor cells and express TCRs that are sufficient to recognize tumor cell-expressed antigens is not clear. It is also not clear if the TCRs selected in an antigen-agnostic manner are patient-specific and can be transferred to allogenic T cells and used for TCR-therapy applications. Almost all current TCR therapy efforts use TCRs targeting known-antigens such as NY-ESO-1 with limited effectiveness (*19*), hence the ability to identify tumor-targeting TCRs in an unbiased manner will be of significant benefit. Here we report a co-culture platform that can achieve a high degree of clonal expansion of memory T cells from peripheral blood mononuclear cells (PBMC), and that can be used to identify tumor-targeting T cell receptors. We further demonstrate that TCRs identified in tumor-targeting TCRs are sufficient to elicit patient-specific tumor-recognition when expressed in allogeneic T cells.

We generated tumor organoids from patients with pancreatic cancer (Extended Data Table 1a) and used it to co-culture with autologous PBMCs. Media containing different ratios of growth factor and cytokines were screened to optimize conditions that support the growth of pancreatic tumor epithelia and PBMC-derived T cells. The conditions were designed to not include extra CD28 agonist (signal 2) to prevent education of naïve T cells to recognize self or non-specific antigens *in vitro* (*20*), and promote expansion of memory T cells that were educated *in vivo* to recognize tumor antigens. Two weeks of co-culture resulted in the complete killing of the tumor organoids and generation of organoid-primed T (opT) cells (Fig. 1a). Once generated, opT cells killed autologous organoids efficiently within 24-48 hours when re-stimulated, demonstrating their cytotoxic activity (Fig. 1c, Extended Data Fig. 1a). The total number of opT cells varied among the three patients studied. Despite starting the co-culture with 0.3 million PBMCs, Pt10 co-culture yielded 3.0 million opT cells, whereas Pt3 co-culture yielded 216 million opT cells (Extended Data Table 1b). To investigate the broader utility of this platform, we explored the ability of this co-culture platform to generate opT cells for tumors generated in genetically-engineered mouse models (GEMM) of breast cancer. Organoids generated from mammary tumors from BRCA/TP53 tumor models were exposed to autologous splenocytes expanded from tumor-bearing animals. These murine opT cells were effective in killing autologous tumors but lacked the ability to kill normal mammary epithelial organoids generated from syngenic mice, demonstrating tumor selectivity for opT cell-mediated killing and generation of IFNg (Extended Data Fig. 1b, c).

**Fig. 1.**
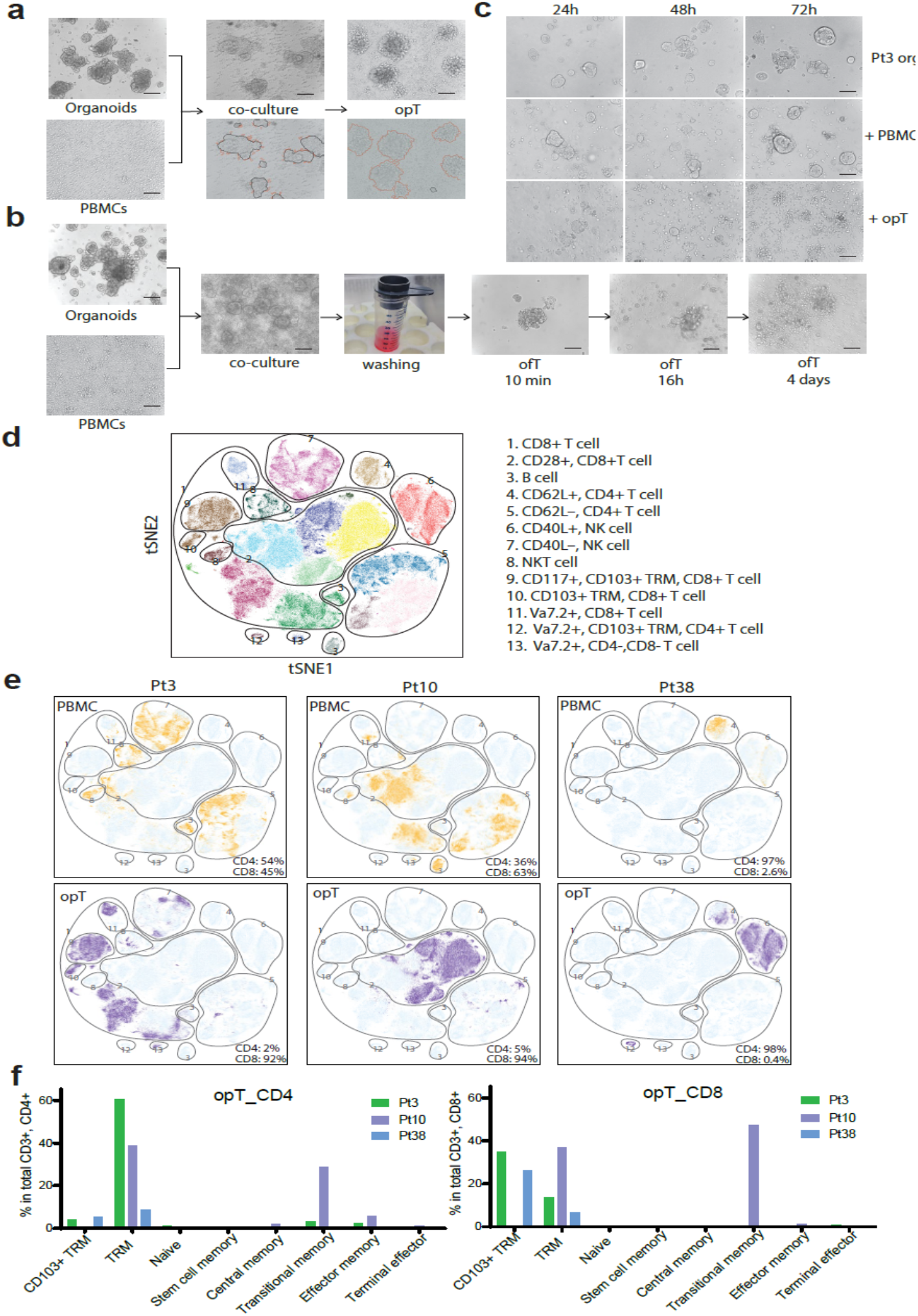
Generation and characterization of tumor-targeting cytotoxic T cells using autologous tumor organoids. **a,** Patient tumor organoids co-cultured with autologous PBMC for 14 days for generation of organoid-primed T (opT) cells. The images are annotated as tumor organoids in “Black” and the T cells located near organoids or clusters of activated T cells in “Red”. **b,** Organoids free of Matrigel were incubated with autologous PBMC for 1 hour and filtered through a 15 μm cell strainer. Organoid and bound T cells were released from the strainer and cultured for 14 days to generate the organoid-fished T (ofT) cells. **c,** Representative phase-contrast images for Pt3 tumor organoids alone or co-culture with autologous PBMC or opT at different time points to demonstrate cell killing. **d,** Annotated image of all possible subtypes from all six samples analyzed. t-SNE plot showing all possible clusters detectable by CyTOF analysis. The CyTOF clusters were assembled into 13 phenotypic groups for interpretation. **e,** Comparison of PBMC (yellow) and opT (purple) cells for the presence of CyTOF clusters against the background of all possible clusters (light blue). The percentage of CD4+ and CD8+ cells within the CD3+ population is shown in the bottom right. **f,** Phenotypes in CD4+ or CD8+ opT cell populations grouped per marker expression. See Table S2 for the makers used to define the phenotypes.

Since the establishment of well growing organoids cultures is not always feasible, in part due to short term survival or slow growth of tumor cells, we explored options to use limited cancer cells to identify tumor-cell-targeting T cells. A high-affinity antigen-TCR interaction is reinforced by the interaction between integrins such as lymphocyte function-associated molecule-1 (LFA-1) on T cells and the intercellular adhesion molecule-1 (ICAM-1) on the tumor cells (*27, 22*). We exploited this biological property and investigated if organoids can be used to purify tumor-targeting T cells in PBMC by an affinity-based purification method. Organoids were incubated with PBMCs for 60 minutes, and unbound immune cells were washed away. The organoid-bound T cells were cultured to permit the killing of the bound tumors cells and expansion of the T cell population (Fig. 1b). These affinity-enriched T cells, herein referred to as organoid-fished T cells (ofT), were effective in killing tumor cells upon re-exposure to autologous organoids, demonstrating their cytotoxic activity as observed for the opT cells. This ofT platform offers opportunities for rapid identification of tumor-targeting T cells from PBMC with limited, transiently cultured, tumor cells. Thus, both opT and ofT platforms have the potential to be broadly applied to generate tumor-targeting T cells for multiple cancers.

To understand the cell types present in human PBMCs and opT cells, we monitored the expression levels of 28 cell surface proteins at a single-cell level by mass cytometry time of flight (CyTOF) analysis (Extended Data Table 1c). CyTOF results from Patient 3 (Pt3), Pt10, and Pt38, were analyzed using t-distributed Stochastic Neighbor Embedding (t-SNE) algorithm to find groups or clusters of cells with similar expression levels of the 28 markers. The clusters were displayed as a t-SNE plot (Fig. 1d), where each dot represents a cell, and the proximity of cells corresponds to the similarity of their protein expression patterns. The analysis of three pairs of matched PBMC and opT cells identified multiple expression clusters, which we divided into 13 subgroups representing distinct cell types and lineages (Fig. 1d). PBMCs from all three patients clustered into multiple phenotypic groups, dominated by T cells (CD4 or CD8) with minor populations of B cell and NK cell types (Fig. 1e).

Next we analyzed how opT cells from the different patients compare with each other. Interestingly, Pt3 and Pt10 opT cells had restricted diversity comprising primarily (89 - 90%) of CD3+ T cells (Fig. 1e). Among the CD3+ cells, Pt3 opT were primarily CD8+ cells whereas, Pt10 had 37% CD4+ and 62% CD8+ cell populations (Fig. 1e). Pt38 opTs, however, had only 33.8% CD3+ T cells (Fig. 1e), which were primarily CD4+. The CD3-cells expressed NK cell markers (Fig. 1e). All the opT cell populations lacked B cells. Thus we observed patient-specific differences in the opT cell phenotype. The differences may not related to specific tumor characteristics because all three patients had stage IV metastatic disease with mutations in KRAS and TP53 at the time of diagnosis. Despite the presence of Grade 2 disease, Pt38 had the shortest survival time after diagnosis (6 months) (Extended Data Table 1a). Although it is possible that the absence of tumor-targeting CD8+ cells in the circulation relates to the poor clinical outcome for Pt38, further studies are need to test this possibility.

OpT cells from all three patients expressed markers of tissue-resident memory T cells (TRM) or CD103+ TRM (Fig. 1f, Extended Data Table 2a). Although there was patient-to-patient variation in the type and percentage of cells with memory phenotype in opT cells, they all had a very low abundance of T cells with naïve or exhausted phenotypes. Furthermore, the patient-to-patient differences in cells with memory phenotype also demonstrate that the culture condition does not normalize the differentiation status of opT cells and permits retention of patient-to-patient variations. TRM phenotype is associated with T cells in tissue bed are considered essential for permanent solid tumor immunity (*23, 24*). Thus, it is likely that our organoid-PBMC co-culture conditions promote the generation of tumor-targeting T cells with the TRM phenotype. Thus our platform represent a unique opportunity for future use of opT and ofT cells in ACT applications in the clinic.

To better understand the biological properties of opT cells, we assessed how opT and unselected PBMCs differ in their ability to respond to autologous tumor cells. To determine changes in the capacity of tumor cells to stimulate T cell proliferation, we monitored cell division using carboxyfluorescein succinimidyl ester (CFSE) labeled PBMC and opT cells cultured in the presence or absence of autologous tumor organoids for four days. PBMC cells showed no significant increase in dividing cell populations as determined by flow cytometry analysis. In contrast, opT cells showed a 15 or 89% increase in the CFSE-low population of cells in Pt3 and Pt38, respectively (Fig. 2a). The ofT cells from Pt10 also showed an increase of 41% in the proliferating cells upon exposure to autologous tumors (Extended Data Fig. 2a, b), demonstrating that both opT and ofT cells respond to autologous tumor organoids by entering the cell cycle.

**Fig. 2.**
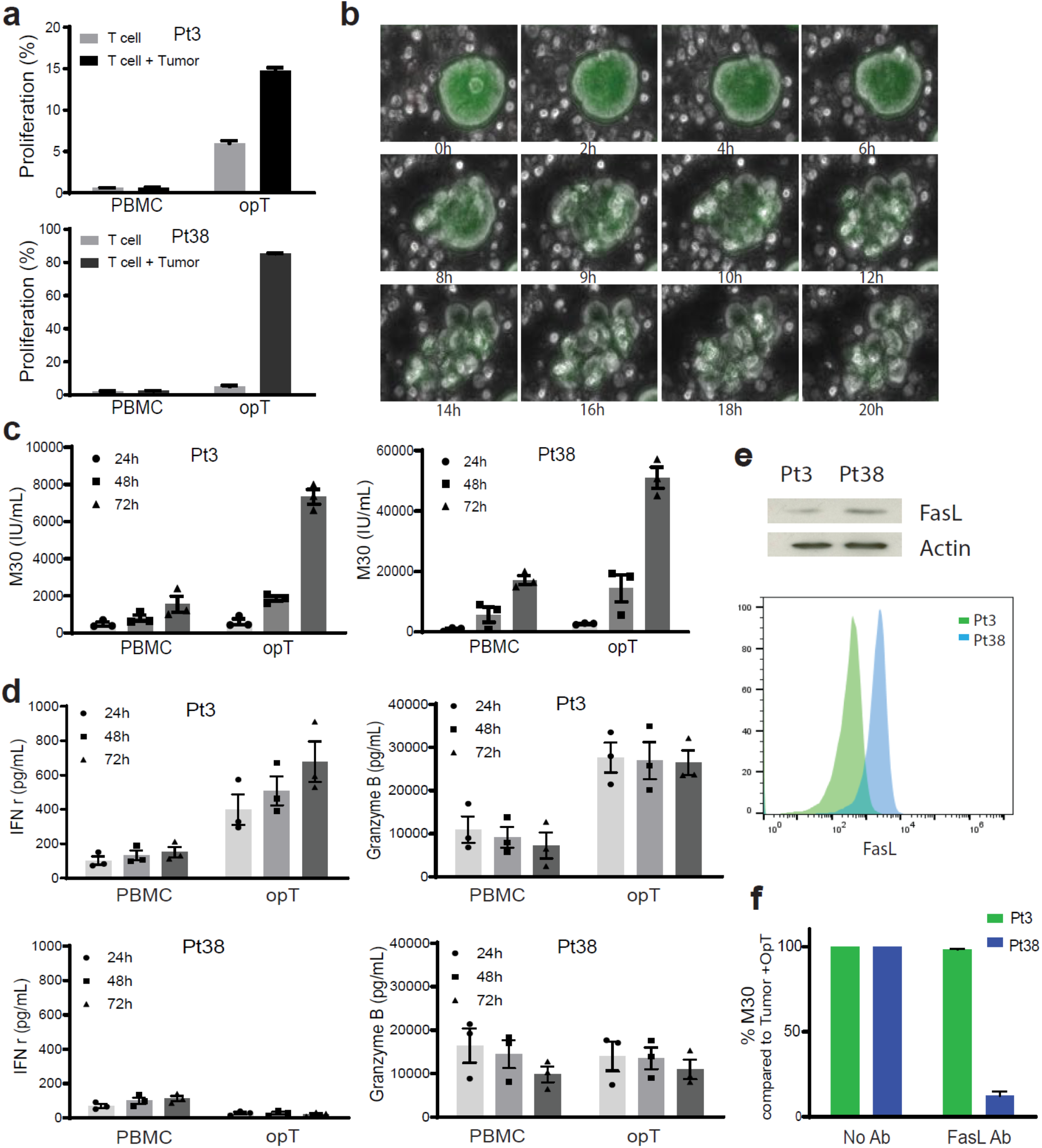
Cytotoxic activity of opT cells. **a,** CFSE (carboxyfluorescein diacetate) labelled PBMC and opT cell proliferation were co-cultured with autologous tumor organoids for four days and changes in percentage of CFSE-low T cell population shown. Mean ± SEM from three independent experiments shown. **b,** Timelapse image of fluorescently labelled organoids (green) incubated with unlabelled opT cells over a period of 20 hours. **c,** Changes in levels of epithelial cell–specific caspase-cleaved cytokeratin 18 (CK18) fragments containing the CK18Asp396 (‘M30’) neo-epitope in the media, quantitated by ELISA. Mean ± SEM from three independent experiments shown. **d,** Interferon gamma and granzyme B produced by PBMC or opT in the presence of autologous tumor organoids at different time points for Pt3 and Pt38. Mean ± SEM from three independent experiments shown. Each dot represents the mean of at least two technical replicates from independent experiments. **e,** FasL expression in opT cells from Pt3 and Pt38 by immublot or flow cytometry. **f,** Relative M30 production from the supernatants of Pt38 or Pt3 co-cultured in the presence or absence of anti-FasL blocking antibody for 72 hours.

To test the ability of opT cells to kill tumor cells, we monitored cell death by live image analysis and performed ELISA to detect epithelial cell-specific caspase-cleaved cytokeratin 18 (CK18) fragments containing the CK18Asp396 (‘M30’) neo-epitope. Live-cell imaging showed that opT cells made contact with organoids starting around 2.0 hours and the initiated killing by 8.0 hours (Fig. 2b). Changes in M30 concentrations in the co-culture media demonstrated that opT cells were 3-5 fold more effective than matched PBMC co-cultures after 48 and 72 hours (Fig. 2c). In addition to opT cells, ofT cells from Pt10 were also effective in killing autologous tumor organoids (Extended Data Fig. 2c).

The Pt3 opT cells with predominantly CD8+ cells induced a greater increase in both IFNγ and granzyme B levels when compared to PBMC in the presence of autologous tumor organoids, demonstrating the ability of Pt3 opT to initiate a cytotoxic response (Fig. 2d). Interestingly, the Pt38 opT cells, showed no difference for granzyme B secretion between PBMC and opT cells and a decrease in IFNγ levels (Fig. 2d), suggesting that Pt38 T cells kill tumor cells by alternate mechanisms. To rule out the possibility that the lack of IFNγ secretion is because of the presence of NK cells, we incubated the CD3+ opT cells with organoids and did not observe a detectable increase in IFNγ (Extended Data Fig. 2d). Consistent with the presence of CD4+ T cell population in Pt38 opT cells, the Pt38 tumor cells express HLA class II proteins (Fig. 3b), suggesting that the tumor cells presented the antigens for CD4+ T cells. Cytotoxic CD4+ cells are known to kill tumor cells by Fas-Fas ligand (FasL) mediated mechanisms (*25, 26*). To understand how the CD3+ T cells in Pt38 opT cells kill tumors, we assessed the FasL expression levels in both Pt3 and Pt38 opT cells and found the Pt38 opT cells had higher levels of expression of FasL than Pt3 (Fig. 2e). Consistently, the anti-FasL antibody significantly inhibited opT cell-mediated killing of Pt38 opT cells with no effect for Pt3 opT cells, demonstrating that Pt38 opT kills tumors using the Fas-FasL pathway whereas Pt3 opT cells kill by IFNγ and granzyme B secretion (Fig. 2f).

**Fig. 3.**
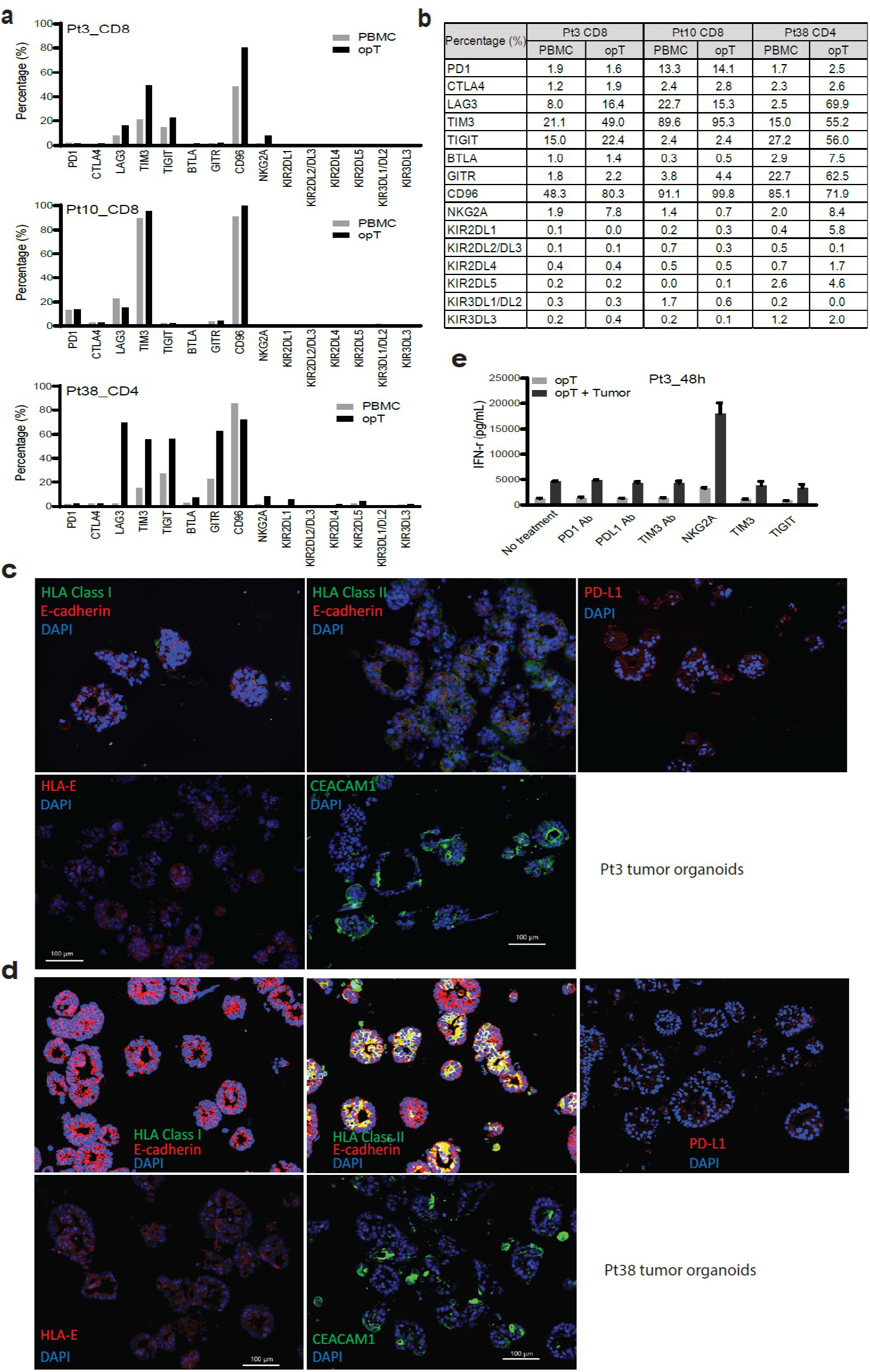
Expression of immunomodulatory proteins and response to checkpoint inhibition in opT cells. **a**, Expression of check-point proteins in opT cells compared to the matched PBMCs by flow cytometry and CyTOF. **b,** Immuno-staining for MHC-I, MHC-II, PD-L1, HLA-E and CEACAM1 in tumor organoids from Pt3 and Pt38. **c,** Changes in IFNγ secretion by opT cells after 48h of pretreatment with anti-PD1, PDL1 and TIM3 blocking antibodies or NKG2A, TIM3, TIGIT and LAG3 protein, in the presence or absence of autologous tumor organoids from Pt3.

Previous studies have demonstrated that in addition to the engagement of TCR with the antigen on tumor cells, T cells are regulated by co-stimulation of receptors and ligands that can either stimulate or inhibit their function (*27*). These co-stimulatory or co-inhibitory (also known as check-point inhibitors) are required for maintaining immune homeostasis in normal tissues (*27*). In cancer tissues, an increase in expression of check-point inhibitors is frequently used for immune evasion, and blockade of these pathways is an established strategy for cancer immunotherapy (*28*). To understand the expression of these markers in opT cells, we monitored levels of 17 immune checkpoint-inhibitors, including the frequently studied PD-1 and CTLA-4, and TIM-3, TIGIT, CD96, LAG-3, BTLA, GITR, NKG2A and KIRs (KIR2DL1-5 and KIR3DL1-3) (*28*) in both PBMC and opT cell populations. We found that there were noticeable inter-patient variations in the expression of check-point inhibitors, and some of the check-point proteins were highly expressed in both PBMC and opT cells. Pt3 opT cells showed increased expression of LAG3, TIM3, TGIT, CD96, and NKG2A; Pt10 opT had limited changes in expression of all check-point proteins tested, and Pt38 opT cells showed an increased expression of LAG3, TIM3, TIGIT, BTLA, GITR, NKG2A, and KIR2DL1 (Fig. 3a). Although NKG2A is primarily expressed in NK cells, many recent studies demonstrated that NKG2A is also expressed by CD8+ T cells within the tumor environment (*18*). To further understand the changes in expression levels of these inhibitory receptors in opT, we monitored changes in expression levels of ligands for the immunomodulatory receptors on tumor cells. HLA class II (ligand for LAG3), PD-L1 (ligand for PD1), HLA-E (ligand for NKG2A), and CEACAM1 (ligand for TIM3) were all expressed on tumor organoids (Fig. 3b, Extended Data Fig. 3a) suggesting that the opT cells were likely to be regulated by these check-point proteins. To determine the functional significance of check-point protein expression, we used inhibitory antibodies or soluble proteins to block PD1, PD-L1, TIM3, TIGIT, LAG3, and NKG2A receptors and found that the NKG2A blockade showed the most potent increase in the IFNγ production both 24 and 48 hours after stimulation (Fig. 3c, Extended Data Fig. 3b) compared to blocking PD1-PDL1 or TIM3 or TIGIT or LAG3. Sorted populations of CD3+ opT cells retained the ability to respond to NKG2A blockade (Extended Data Fig. 3c, d), demonstrating that NKG2A blockade functions as a promoter of cytotoxic T cell activity and not due to the NK cells present in Pt3 opT cells. Our observations are consistent with a report that identifies NKG2A as an important immune regulatory protein for tumor-specific T cells, in addition to its role in NK cells(*29*). The results also demonstrate the ability to use this co-culture platform to identify personalized check-point inhibition strategy for effective immunotherapy.

To investigate if opT cells are an activated polyclonal population of the T cells or clonally expanded populations of tumor-targeting T cells, we sequenced TCR for matched PBMC and opT cells from Pt3, Pt10, and Pt38. More than 150,000 TCR β-chains were sequenced from PBMC and opT cells. As expected, in PBMC from both Pt3 and Pt10, no TCR was represented more than 3.0%, demonstrating a polyclonal nature of the population (Fig. 4a). However, Pt3 opT cells were oligoclonal with one dominant TCR (referred to as Organoid Selected T cell Receptor-1, OSR-1), representing 81% of the opT cell population. OSR1 was present in only 2.3% of PBMC, thus representing a more than 35 fold clonal expansion after the co-culture with autologous tumors. Four other clones contributed to an additional 18 percent for the Pt3 opT cell population (Fig. 4a, b). In Pt38, the T cell clone OSR11 was undetectable in PBMC but was enriched to 90.4% in opT, demonstrating a more than 270,000 fold clonal expansion after the coculture. Similar results were also observed for ofT cells, where OSR16 was detected in less than 0.0005% TCRs in PBMC but increased to 33.6% in ofT for Pt10 (Extended Data Fig. 4b) representing more than 63,000 fold clonal expansion. Thus, in each of the three opT cell populations, five TCRs made-up 89.8% to 99.0% of the diversity in the >150K TCRs analyzed, demonstrating an unexpected and powerful clonal expansion, which can serve as a platform for empirical identification of tumor-targeting T cell clones from peripheral blood of patients with pancreas cancer. Furthermore, all the fifteen highly expanded TCRs were distinct between these three opT cell populations, demonstrating antigen diversity between these patients. The lack of common TCRs between opT populations emphasizes the opportunity for personalizing immunotherapy approaches, and also rules out non-specific clonal selection induced by shared antigens present in our cell culture conditions.

**Fig. 4.**
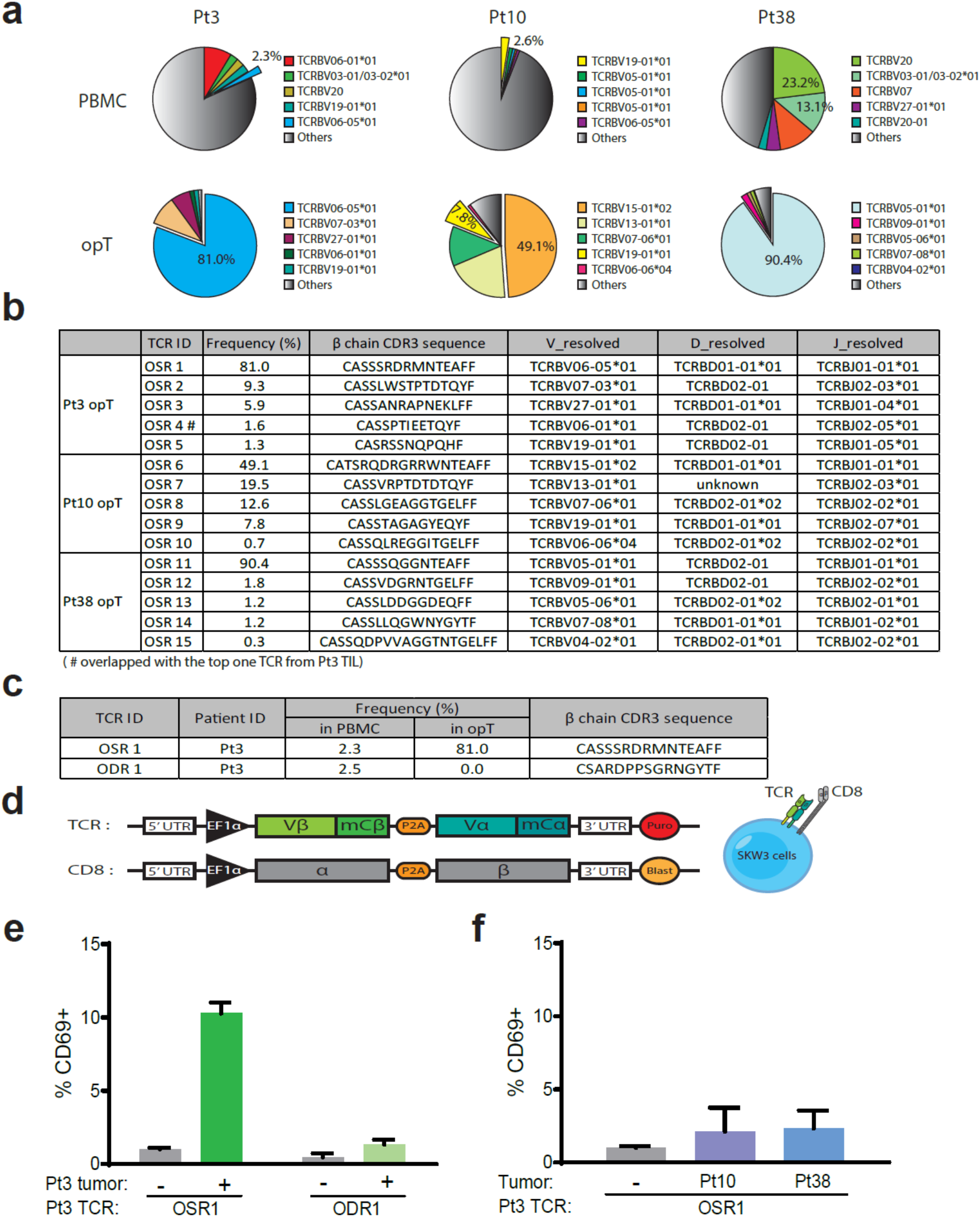
Clonal expansion in opT cells and identification of TCR. **a,** The variable beta-chain of more than 150,000 independent T cell receptors were sequenced for PBMC and opT cells and the relative representation of a given TCRs as percentage of the total is shown in pie charts. **b,** Frequencies and CDR3 sequences for the top five organoid-selected TCR (OSR from each patient. **c,** TCR enriched in Pt3 opT and organoid-depleted TCR (ODR1) used for generation of recombinant TCR and CDR3β sequence. **d,** The CDR3 regions were used to generate a chimeric TCR that comprised of human Vα and Vβ chains and mouse constant α and β chains and coexpressed with human CD8 gene in SKW-3 cells. **e,** Expression of T cell activation marker (CD69) in TCR-expressing SKW-3 cells exposed to autologous (Pt3) tumor organoids. **f,** Expression of CD69 in OSR1 expressing SKW-3 cells exposed to allogenic (Pt10 and Pt38) tumor organoids.

We compared the β chain of top 5 TCRs from the opT cells and TlLs with a TCR database from 120 healthy individuals to determine if the TCRs expressed by opT cells are present in health individuals. One TCR from Pt38 opT cells was present in 28 healthy individuals with a highest frequency of 1×10-5 6/15, five other TCRs were present in less than 10 individuals and nine were not observed in any of the 120 healthy individuals (Extended Data Table 2c). We next analyzed if the TCRs identified in opT are patient-specific. Interestingly, TCRs identified in one patient was not present in the PBMC of the other PDAC patients suggesting that there may be significant patient-specificity(Extended Data Table 2d). However, a larger cohort of opT TCRs need to analyzed before arriving at any broad conclusion.

To understand how the TCRs selected in our co-culture platform compares with the T cells naturally enriched in the tumor bed *in vivo*, we compared the TCRs identified in expanded TILs from Pt3 tumor with Pt3 opT cells. There was a 50% overlap among the top 10 TCRs between TILs and opT cells (Extended Data Table 2b), including OSR4 (Fig. 4b), suggesting that there are similarities and differences in the clonal selection that occurs in the tumor bed and the coculture platform. Since the co-culture lacks inhibitory factors contributed by the microenvironment, it likely creates an opportunity for unrestricted T cell-tumor cell interactions.

Next, we investigated if the TCRs present in opT cells retain the ability to recognize tumor cells when transferred to un-trained T cells. We determined the sequence of both alpha and beta chain of the TCRs in opT cells and selected the top five TCRs to generate a chimeric TCR (Fig. 4b). TCR sequences that were detectable in PBMC but lost during the opT selection process (Organoid Depleted T cell receptors (ODR)) were used as negative controls (Fig. 4c). The chimeric receptor was expressed in SKW-3, a T cell line that lacks endogenous TCR(*30*) to generate populations of SKW-3 cells expressing recombinant TCRs identified in opT cells. The SKW-3 cells were exposed to organoids and monitored for changes in the expression of the T cell activation marker CD69. The SKW-3 cells expressing the positively-selected TCRs induced a significant increase in the expression of CD69 upon co-culture with autologous tumors (Fig. 4e). The TCR-expressing SKW-3 cells did not respond to exposure to allogenic tumor organoids (Fig. 4f), highlighting the presence of patient-specific antigens that are recognized by the TCRs selected in opT cell populations. To determine if the TCRs in the CD4+ opT cells of Pt38 also retained the ability to recognize tumors in SKW-3 cells, we sequenced the alpha chains in Pt38 opT population. Interestingly, unlike the OSR11 beta chain that constituted 90.4% in opT TCRs, we observed two alpha chains at 46% each, suggesting that the the OSR11 beta chain may exist as two different TCR pairs in Pt38 opT cells. We generated two chimeric TCRs that share the same beta chain but differ in the alpha chain and expressd them in SKW-3 cells. Cells expressing OSR11-2, but not OSR11-1, increased the expression of CD69 upon co-culture with Pt38 organoids, demonstating that only one of the two α/β pair was functional in recognizing antigens on Pt38 tumor cells (Extended Data Fig. 4c).Thus, we demonstrate that the TCRs selected in CD8+ or CD4+ opT cells was sufficient to confer tumor-targeting ability in a patient-specific manner.

Solid tumors have a heterogeneous pattern of tumor-associated antigen expression, which is thought to contribute to immune escape and decrease the efficacy of immune therapy. Identifying tumor-selective T cell receptors with limited or no cross-reactivity to normal cells will be a significant advance towards the identification of new immune therapy approaches for solid tumors. Here we report the development of an autologous pancreas tumor organoid-immune cell culture platform and demonstrate that the tumor-targeting T cells with a memory phenotype can be isolated from the patient’s peripheral blood and that these T cells undergo clonal expansion expressing TCRs that are tumor and patient-selective. Our findings not only open new opportunities for generating tumor-targeting T cells for personalized adoptive cell therapy opportunities, but also for using this co-culture as a platform for designing a personalized checkpoint inhibitor combination immunotherapies for pancreatic cancer patients.

## Acknowledgments

We thank Dr. Kai Wucherpfenning for critical input on the manuscript and members of the Muthuswamy and Hidalgo laboratories for their helpful and critical suggestions during the course of the study.

## Author contributions

MH and SKM conceived the study. SKM, QM, SX, and MH designed the experiments. QM and SX performed the experiments and prepared the illustrations. GKG anlalyzed CyTOF data and made illustrations. MHD and SE assisted with TCR vectors and SKW-3 analysis. LH and DA generated the patient tumor organoid lines. WL assisted with SKW-3 experiments and EF contributed to immunofluorescence. SKM recruited the collborators and SKM and QM interpreted all the results. SKM, QM, SX, and MH co-wrote the manuscript.

---

**Extended Data Fig. 1.**
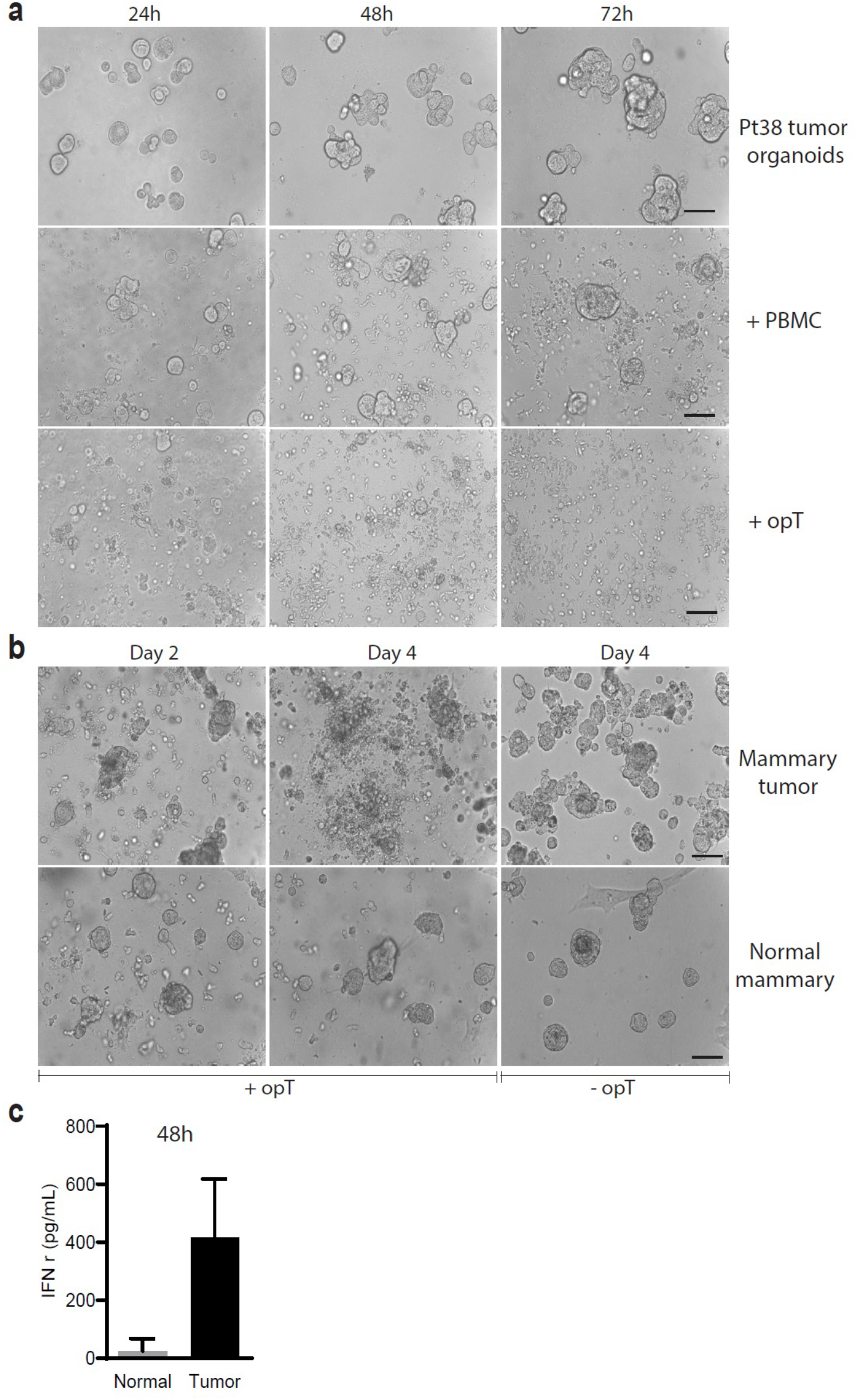
**a,** Representative phase-contrast images for pt38 tumor organoids alone or coculturewith autologous PBMC or opT at different time points to demonstrate cell killing. Bar: 50μm. **b,** Representative phase-contrast images of mouse mammary tumor organoid orsyngenic normal mammary epithelial oragnoids incubated with or without opT cells for indicated times. Bar: 50μm. **c,** Media supernatant from mouse organoids co-cultured for two days with opT cells analyzed for IFNg levels by ELISA. **d,** Phenotype of T cell populations in PBMC as analyzed by CyTOF.

**Extended Data Fig. 2.**
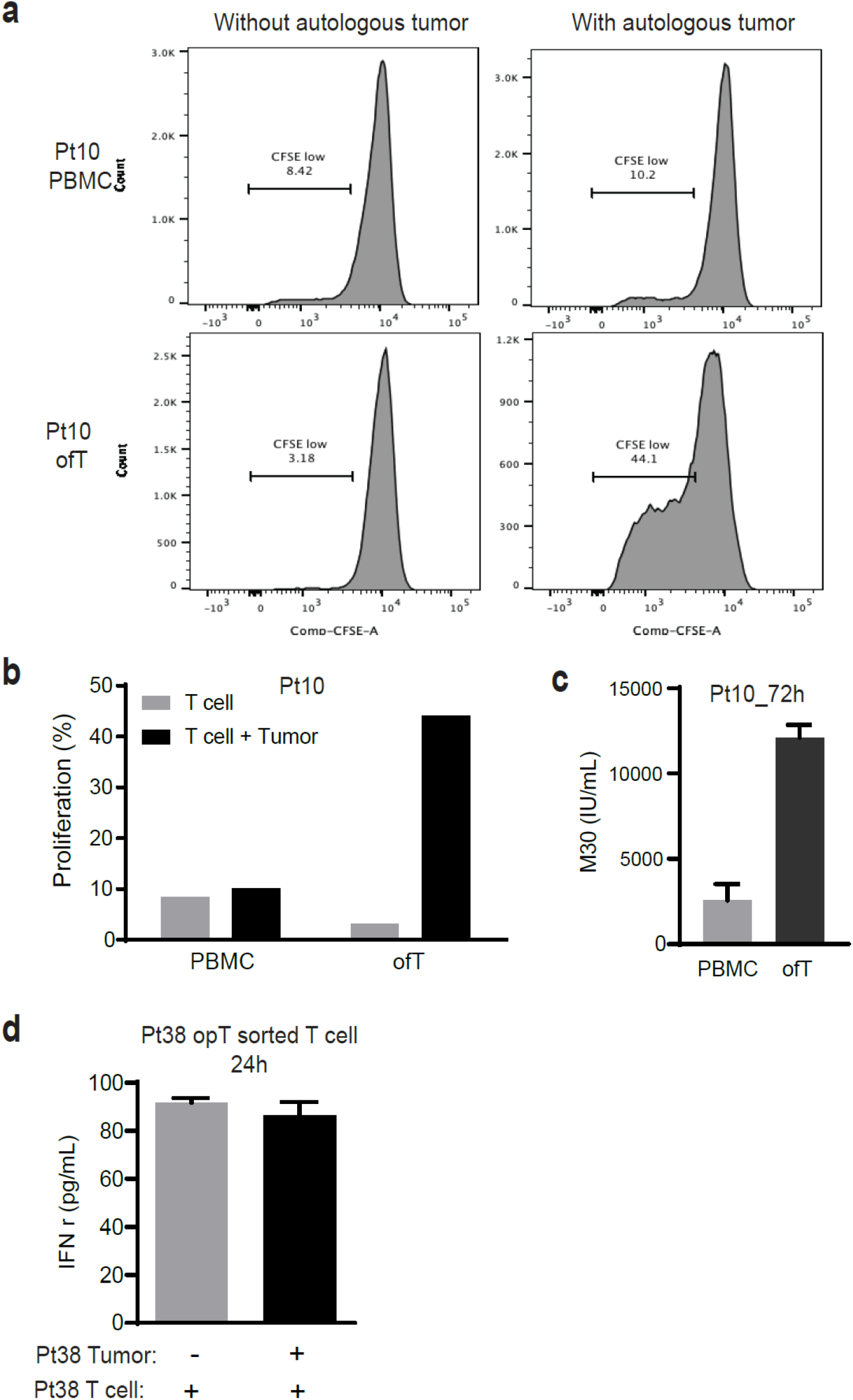
**a,** CFSE labelled PBMC or ofT cells from pt10 were incubated with autologous tumor organoids and analyzed by flow cytometry after four days. Percentage of CFSE-low population shown. **b,** Graphical representation of the changes in CFSE-low populations shown in panel A. **c,** PBMC or ofT cells from pt10 were incubated with autologous organoids for 72 hours and media supernatant analyzed for changes in levels of the M30, the caspase-cleave CK18 marker to monitor cell death. **d,** CD3+ pt38 opT cells were incubated with or without autologous tumor organoids and media analyzed for IFNg levels after 24 hours.

**Extended Data Fig. 3.**
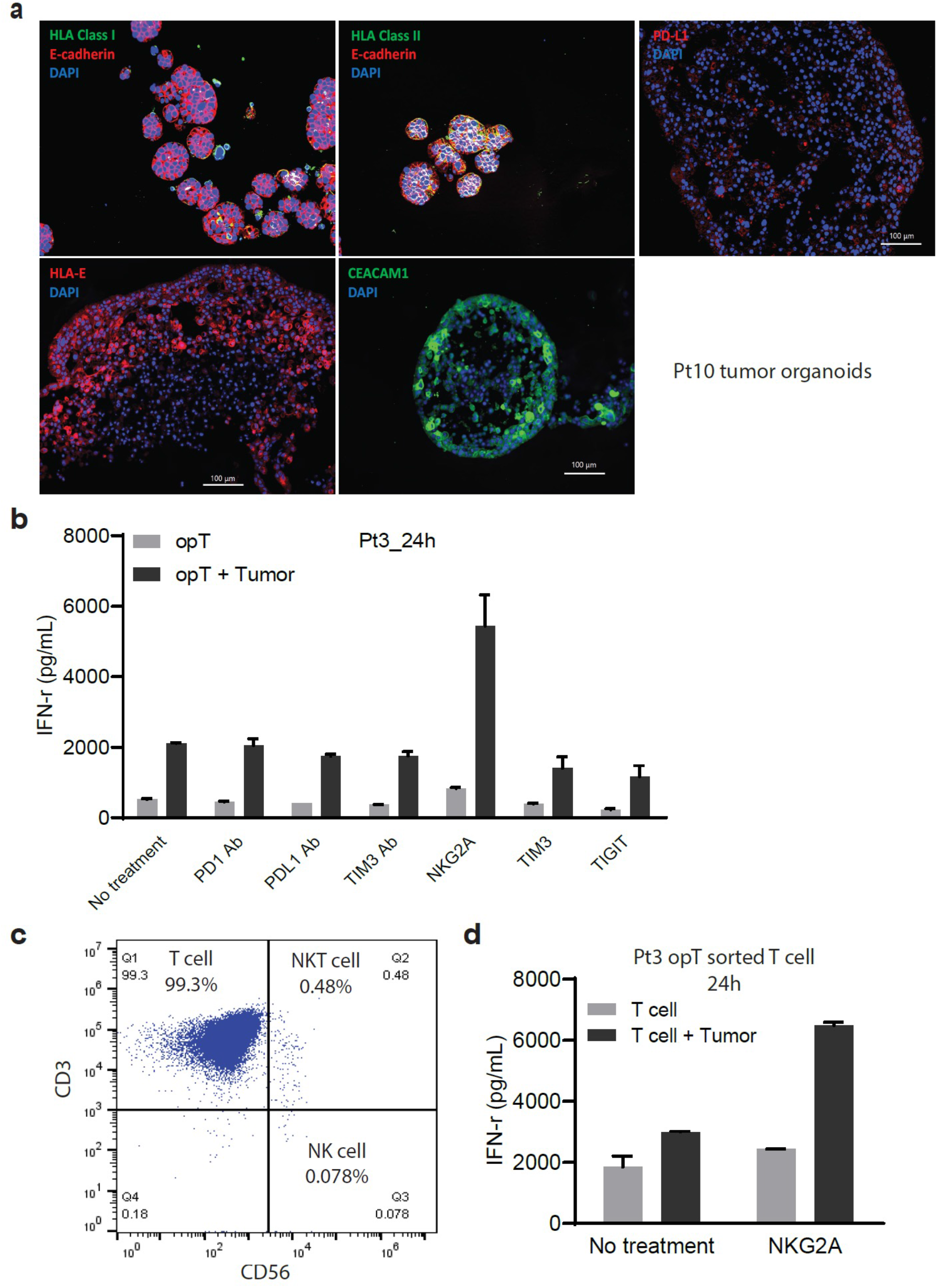
**a,** Immuno-staining for MHC-I, MHC-II, PD-L1, HLA-E and CEACAM1 in tumor organoids from pt10. **b,** Changes in IFNg secretion by opT cells after 24h of pretreatment with anti-PD1, PDL1 and TIM3 blocking antibodies or NKG2A, TIM3, TIGIT and LAG3 protein, in the presence or absence of autologous tumor organoids from pt3. **c,** opT cells from pt3 sorted for T cell (CD3) and NK cell (CD56) markers. **d,** CD3+ opT cells from pt3 were incubated with autologous tumor organoids in the presence or absence of NKG2A for 24 hours and media analyzed for changes in IFNg levels.

**Extended Data Fig. 4.**
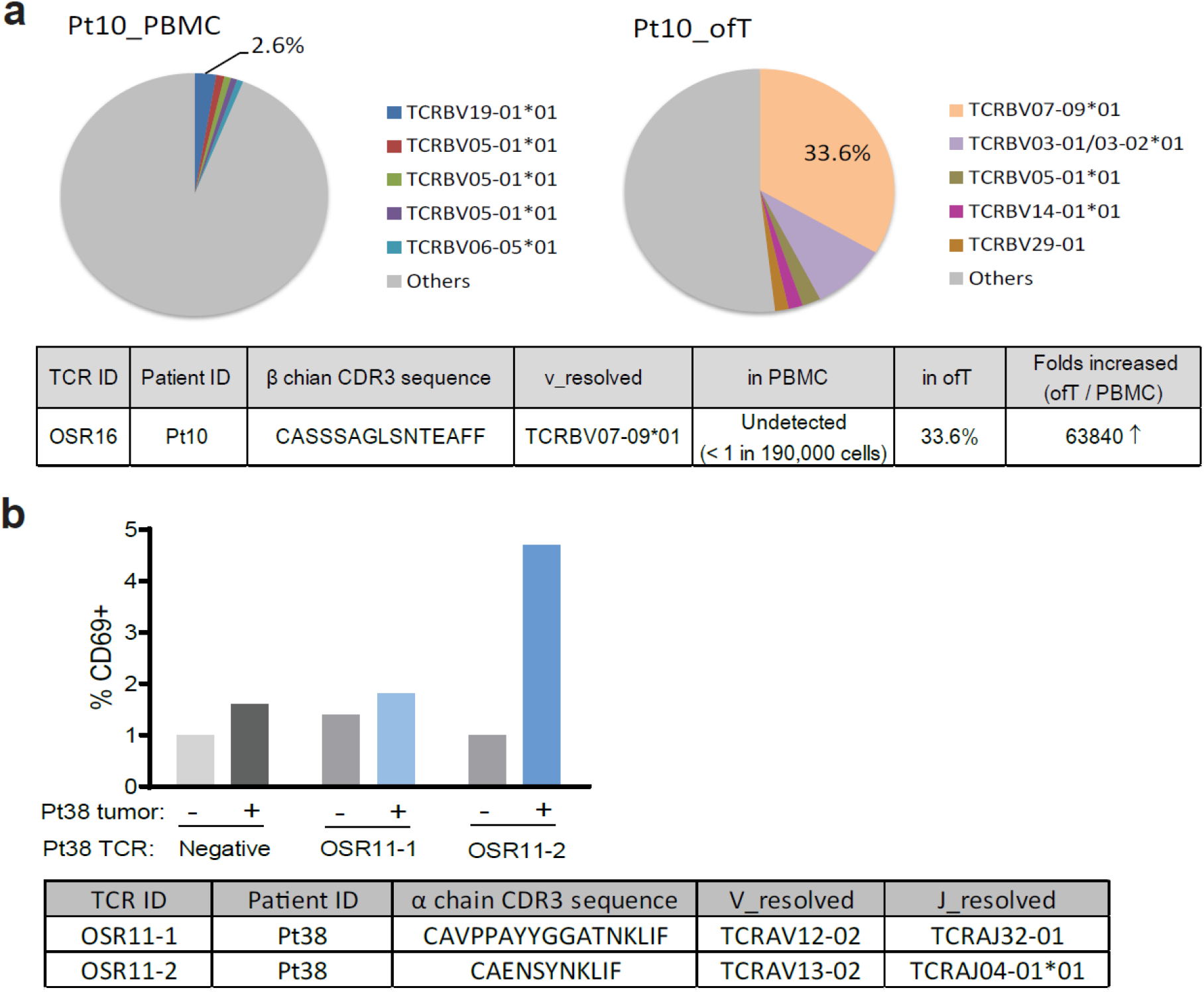
**a,** The variable beta-chain of more than 150,000 independent T cell receptors were sequenced for PBMC and ofT cells of pt10 and the relative representation of a given TCRs as percentage of the total is shown in pie charts. The dominant TCR identified in ofT (OSR16) is shown in the table below. **b,** The beta chain of OSR11 that from pt38 that was represented at 90.4% was matched with two equally represented alpha chains (46% each) to generate two independent TCRs (OSR11-1 and ISR11-2) that share the beta chain. The chimera were expressed in SKW-3 and exposed to pt38 tumor organoids and changes in expression of CD69 were monitored by flow cytometry. SKW-3 cells expressing CD8 alone and not chimeric TCR was used as negative control.

**Extended Data Table 1.**
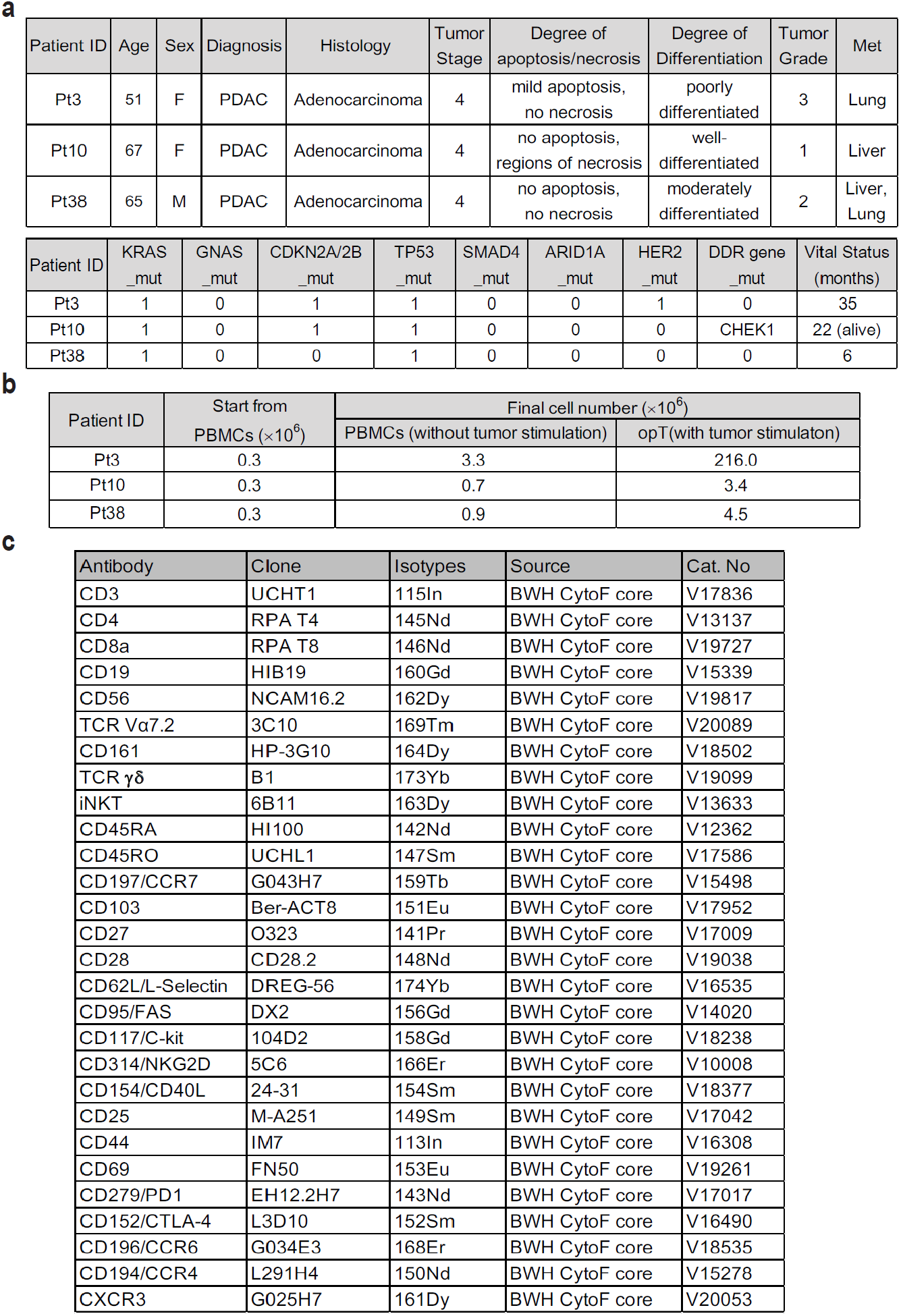
**a,** Information pertaining to the three PDAC patient used in this study. **b,** Changes in cell number in response to organoid co-culture. **c,** Antibodies used for CyTOF analysis.

| Patient ID | Age | Sex | Diagnosis | Histology | Tumor Stage | Degree of apoptosis/necrosis | Degree of Differentiation | Tumor Grade | Met |
| --- | --- | --- | --- | --- | --- | --- | --- | --- | --- |
| Pt3 | 51 | F | PDAC | Adenocarcinoma | 4 | mild apoptosis, no necrosis | poorly differentiated | 3 | Lung |
| Pt10 | 67 | F | PDAC | Adenocarcinoma | 4 | no apoptosis, regions of necrosis | well-differentiated | 1 | Liver |
| Pt38 | 65 | M | PDAC | Adenocarcinoma | 4 | no apoptosis, no necrosis | moderately differentiated | 2 | Liver, Lung |

| Patient ID | KRAS _mut | GNAS _mut | CDKN2A/2B _mut | TP53 _mut | SMAD4 _mut | ARID1A _mut | HER2 _mut | DDR gene _mut | Vital Status (months) |
| --- | --- | --- | --- | --- | --- | --- | --- | --- | --- |
| Pt3 | 1 | 0 | 1 | 1 | 0 | 0 | 1 | 0 | 35 |
| Pt10 | 1 | 0 | 1 | 1 | 0 | 0 | 0 | CHEK1 | 22 (alive) |
| Pt38 | 1 | 0 | 0 | 1 | 0 | 0 | 0 | 0 | 6 |

**b**
| Patient ID | Start from PBMCs ($\times 10^6$) | Final cell number ($\times 10^6$) | |
| --- | --- | --- | --- |
|  |  | PBMCs (without tumor stimulation) | opT (with tumor stimulation) |
| Pt3 | 0.3 | 3.3 | 216.0 |
| Pt10 | 0.3 | 0.7 | 3.4 |
| Pt38 | 0.3 | 0.9 | 4.5 |

| Antibody | Clone | Isotypes | Source | Cat. No |
| --- | --- | --- | --- | --- |
| CD3 | UCHT1 | 115In | BWH CytoF core | V17836 |
| CD4 | RPA T4 | 145Nd | BWH CytoF core | V13137 |
| CD8a | RPA T8 | 146Nd | BWH CytoF core | V19727 |
| CD19 | HIB19 | 160Gd | BWH CytoF core | V15339 |
| CD56 | NCAM16.2 | 162Dy | BWH CytoF core | V19817 |
| TCR Vα7.2 | 3C10 | 169Tm | BWH CytoF core | V20089 |
| CD161 | HP-3G10 | 164Dy | BWH CytoF core | V18502 |
| TCR $\gamma\delta$ | B1 | 173Yb | BWH CytoF core | V19099 |
| iNKT | 6B11 | 163Dy | BWH CytoF core | V13633 |
| CD45RA | HI100 | 142Nd | BWH CytoF core | V12362 |
| CD45RO | UCHL1 | 147Sm | BWH CytoF core | V17586 |
| CD197/CCR7 | G043H7 | 159Tb | BWH CytoF core | V15498 |
| CD103 | Ber-ACT8 | 151Eu | BWH CytoF core | V17952 |
| CD27 | O323 | 141Pr | BWH CytoF core | V17009 |
| CD28 | CD28.2 | 148Nd | BWH CytoF core | V19038 |
| CD62L/L-Selectin | DREG-56 | 174Yb | BWH CytoF core | V16535 |
| CD95/FAS | DX2 | 156Gd | BWH CytoF core | V14020 |
| CD117/C-kit | 104D2 | 158Gd | BWH CytoF core | V18238 |
| CD314/NKG2D | 5C6 | 166Er | BWH CytoF core | V10008 |
| CD154/CD40L | 24-31 | 154Sm | BWH CytoF core | V18377 |
| CD25 | M-A251 | 149Sm | BWH CytoF core | V17042 |
| CD44 | IM7 | 113In | BWH CytoF core | V16308 |
| CD69 | FN50 | 153Eu | BWH CytoF core | V19261 |
| CD279/PD1 | EH12.2H7 | 143Nd | BWH CytoF core | V17017 |
| CD152/CTLA-4 | L3D10 | 152Sm | BWH CytoF core | V16490 |
| CD196/CCR6 | G034E3 | 168Er | BWH CytoF core | V18535 |
| CD194/CCR4 | L291H4 | 150Nd | BWH CytoF core | V15278 |
| CXCR3 | G025H7 | 161Dy | BWH CytoF core | V20053 |

**Extended Data Table 2.** **a,** Markers used as key to define T cell phenotypes. **b,** Comparisons of TCR beta chain sequences identified in opT and TILs from pt 3. **c,** Frequency of the top 5 TCRs from opT, ofT and TIL in the PBMC of 120 healthy donors. **d,** Cross patient comparison of TCRs in opT cells with TCRs identified in PBMCs of other PDAC patients analyzed in this study.

| Phenotype Key |  |
| --- | --- |
| T cell | CD3+ |
| CD4+ or CD8+ |  |
| Tissue-resident memory (TRM) | CD69+,CD103-,CCR7-,CD45RO+ |
| CD103+ TRM | CD69+,CD103+,CCR7-,CD45RO+ |
| Naive | CD69-,CD103-,CD95-,CD28+,CCR7+,CD45RO- |
| Stem Cell Memory | CD69-,CD103-,CD95+,CD28+,CCR7+,CD45RO- |
| Central memory | CD69-,CD103-,CD95+,CD28+,CCR7+,CD45RO+ |
| Transitional memory | CD69-,CD103-,CD95+,CD28+,CCR7-,CD45RO+ |
| Effector memory | CD69-,CD103-,CD95+,CD28-,CCR7-,CD45RO+ |
| Terminal effector | CD69-,CD103-,CD95+,CD28-,CCR7-,CD45RO- |

b
| | TCR Rank | Frequency (%) | $\beta$ chain CDR3 sequence | V_resolved | D_resolved | J_resolved |
| --- | --- | --- | --- | --- | --- | --- |
| Pt3 opT | 1 | 81.0 | CASSSRDRMNTAEFF | TCRBV06-05*01 | TCRBD01-01*01 | TCRBJ01-01*01 |
|  | 2 | 9.3 | CASSLWSTPTDTQYF | TCRBV07-03*01 | TCRBD02-01 | TCRBJ02-03*01 |
|  | 3 | 5.9 | CASSANRAPNEKLFF | TCRBV27-01*01 | TCRBD01-01*01 | TCRBJ01-04*01 |
|  | 4 | 1.6 | CASSPTIETQYF | TCRBV06-01*01 | TCRBD02-01 | TCRBJ02-05*01 |
|  | 5 | 1.3 | CASRSSNQPHF | TCRBV19-01*01 | TCRBD02-01 | TCRBJ01-05*01 |
|  | 6 | 0.20 | CASSLAGEVEDTQYF | TCRBV07-07*01 | TCRBD02-01*02 | TCRBJ02-03*01 |
|  | 7 | 0.13 | CASSTGDEQFF | TCRBV02-01 | TCRBD01-01*01 | TCRBJ02-01*01 |
|  | 8 | 0.11 | CASSEQRGGFNEQFF | TCRBV02-01 | TCRBD02-01*01 | TCRBJ02-01*01 |
|  | 9 | 0.04 | CASSTTLRDRGLANTGELFF | TCRBV28-01*01 | TCRBD01-01*01 | TCRBJ02-02*01 |
|  | 10 | 0.03 | CASSYSDSGKIYYEQYF | TCRBV06-07*01 | TCRBD02-01*02 | TCRBJ02-07*01 |
| Pt3 TIL | 1 | 50.8 | CASSPTIETQYF | TCRBV06-01*01 | TCRBD02-01 | TCRBJ02-05*01 |
|  | 2 | 30.5 | CASSQDGIAGETQYF | TCRBV04-01*01 | TCRBD02-01*01 | TCRBJ02-05*01 |
|  | 3 | 14.1 | CSAADPYSYEQYF | TCRBV20 | TCRBD02-01 | TCRBJ02-07*01 |
|  | 4 | 2.8 | CASSGDSYGYTF | TCRBV12-03/12-04*01 | TCRBD01-01*01 | TCRBJ01-02*01 |
|  | 5 | 1.3 | CASRPSRATNEKLFF | TCRBV27-01*01 | unknown | TCRBJ01-04*01 |
|  | 6 | 0.28 | CATSRDQGLASNEQFF | TCRBV15-01*02 | TCRBD02-01 | TCRBJ02-01*01 |
|  | 7 | 0.04 | CASSEDRRYGYTF | TCRBV07-02 | TCRBD01-01*01 | TCRBJ01-02*01 |
|  | 8 | 0.01 | CAWSVRGLPQHF | TCRBV30-01*01 | TCRBD02-01*01 | TCRBJ01-05*01 |
|  | 9# | 0.01 | CASSRVGLDYEYF | TCRBV21-01*01 | TCRBD02-01 | TCRBJ02-07*01 |
|  | 10 | 0.01 | CASRSSNQPHF | TCRBV19-01*01 | TCRBD02-01 | TCRBJ01-05*01 |

(Same background color: overlapped TCRs between Pt3 opT and TIL; #: overlapped with rank 52 TCR in Pt3 opT) c
|  | TCR ID | TCR Rank | No. of healthy controls with the same TCR in total 120 | Highest TCR frequency |
| --- | --- | --- | --- | --- |
| Pt3 opT | OSR 1 | 1 | 0 | — |
|  | OSR 2 | 2 | 1 | 3.00E-06 |
|  | OSR 3 | 3 | 0 | — |
|  | OSR 4 | 4 | 0 | — |
|  | OSR 5 | 5 | 6 | 4.3569E-06 |
| Pt10 opT | OSR 6 | 1 | 0 | — |
|  | OSR 7 | 2 | 0 | — |
|  | OSR 8 | 3 | 0 | — |
|  | OSR 9 | 4 | 0 | — |
|  | OSR 10 | 5 | 0 | — |
| Pt38 opT | OSR 11 | 1 | 25 | 1.15821E-05 |
|  | OSR 12 | 2 | 8 | 7.2296E-06 |
|  | OSR 13 | 3 | 0 | — |
|  | OSR 14 | 4 | 0 | — |
|  | OSR 15 | 5 | 0 | — |
| Pt10 ofT | OSR 16 | 1 | 0 | — |
|  | OSR 17 | 2 | 0 | — |
|  | OSR 18 | 3 | 7 | 1.66715E-05 |
|  | OSR 19 | 4 | 0 | — |
|  | OSR 20 | 5 | 0 | — |
| Pt3 TIL | TIR 1 | 1 | 0 | — |
|  | TIR 2 | 2 | 0 | — |
|  | TIR 3 | 3 | 2 | 6.1257E-06 |
|  | TIR 4 | 4 | 0 | — |
|  | TIR 5 | 5 | 0 | — |

(—, undetected.) d
|  | TCR ID | TCR Rank | Percentage (%) in |  |  |
| --- | --- | --- | --- | --- | --- |
|  |  |  | Pt3 PBMC | Pt10 PBMC | Pt38 PBMC |
| Pt3 opT | OSR 1 | 1 | 2.3 | — | — |
|  | OSR 2 | 2 | 0.7 | — | — |
|  | OSR 3 | 3 | 0.5 | — | — |
|  | OSR 4 | 4 | 8.6 | — | — |
|  | OSR 5 | 5 | 2.4 | — | — |
| Pt10 opT | OSR 6 | 1 | — | 0.2 | — |
|  | OSR 7 | 2 | — | 0.7 | — |
|  | OSR 8 | 3 | — | — | — |
|  | OSR 9 | 4 | — | 2.6 | — |
|  | OSR 10 | 5 | — | — | — |
| Pt38 opT | OSR 11 | 1 | — | — | — |
|  | OSR 12 | 2 | — | — | 1.7 |
|  | OSR 13 | 3 | — | — | 0.5 |
|  | OSR 14 | 4 | — | — | 0.5 |
|  | OSR 15 | 5 | — | — | 2.0 |

(—, undetected.)

**Extended Data Table 3.** Antibodies used for flow cytometry analysis.

| Antibody | Color | Clone | Company | Catalogue No. |
| --- | --- | --- | --- | --- |
| CD3 | PE | HIT3a | Biolegend | 300308 |
| CD3 | BV605 | UCHT1 | Biolegend | 300460 |
| CD4 | APC | OKT4 | Biolegend | 317416 |
| CD4 | BV650 | OKT4 | Biolegend | 317435 |
| CD8 | FITC | SK1 | BD | 347313 |
| CD8 | PerCP | SK1 | Biolegend | 344707 |
| LAG-3 (CD223) | BV 605 | 11C3C65 | Biolegend | 369323 |
| TIM-3 (CD366) | APC | F38-2E2 | Biolegend | 345011 |
| TIGIT (VSTM3) | APC | A15153G | Biolegend | 372705 |
| BTLA (CD272) | PE | MIH26 | Biolegend | 344505 |
| GITR (CD357) | PerCP/Cyanine5.5 | 108-17 | Biolegend | 371217 |
| CD96 | PE | NK92.39 | Biolegend | 338405 |
| NKG2A | BV711 | 131411 (RUO) | BD | 747919 |
| KIR2DL1 (CD158a) | PE | REA284 | Miltenyi Biotec | 130-120-586 |
| KIR2DL2/DL3 (CD158b) | PerCP | DX27 | Miltenyi Biotec | 130-099-700 |
| KIR2DL4 (CD158d) | FITC | REA768 | Miltenyi Biotec | 130-112-536 |
| KIR2DL5 (CD158f) | PE | REA955 | Miltenyi Biotec | 130-115-911 |
| KIR3DL1/DL2(CD158e/k) | FITC | REA970 | Miltenyi Biotec | 130-116-280 |
| KIR3DL3 (CD158z) | AF647 | 1136B | R&D systems | FAB8919R-025 |

